# Live-cell imaging and mathematical analysis of the “community effect” in apoptosis

**DOI:** 10.1101/2022.07.21.500970

**Authors:** Diane Coursier, David Coulette, Hélène Leman, Emmanuel Grenier, Gabriel Ichim

## Abstract

As a cellular intrinsic mechanism leading to cellular demise, apoptosis was thoroughly characterized from a mechanistic perspective. Nowadays there is an increasing interest in describing the non-cell autonomous or community effects of apoptosis, especially in the context of resistance to cancer treatments. Transitioning from cell-centered to cell population-relevant mechanisms adds a layer of complexity for imaging and analyzing an enormous number of apoptotic events. In addition, the community effect between apoptotic and living cells is difficult to be taken into account for complex analysis. We describe here a robust and easy to implement method to analyze the interactions between cancer cells, while under apoptotic pressure. Using this approach we showed as proof-of-concept that apoptosis is insensitive to cellular density, while the proximity to apoptotic cells increases the probability of a given cell to undergo apoptosis.

## Introduction

Inconspicuous at the single cell level, apoptosis is a powerful mechanism when triggered at the organismal level. It is the sculpting force behind all multicellular organism morphogenesis, ensuring all organs have the right size, supernumerary cells are removed and neurons innervate the correct targets.

In cancer, apoptosis is an efficient guardian against oncogenic transformation and it is the preferred mechanism employed by tumor suppressor proteins to eliminate dangerous rogue cells (*1*). For instance, p53 can trigger apoptosis by up-regulating the expression of several pro-apoptotic proteins (*1*). Apoptosis is undeniably the main executioner of most cancer drugs and radiotherapy (*2*).

Apoptosis is triggered when one of two distinct pathways is engaged: the extrinsic or the intrinsic. Extrinsic apoptosis relies on the stimulation of a death receptor-family member (*e.g.,* TRIAL-R, FAS or TNF-R) by a TNF-related cytokine, and activation of caspase 8 and 10 via the death-inducing signaling complex (DISC) (*3*). The intrinsic pathway is initiated by an intracellular death stimulus such as DNA damage, chemo- and radiotherapy or tumor suppressor activation. In this case, the point of no return is mitochondrial outer-membrane permeabilization (or MOMP) followed by cytochrome *c* release, apoptosome assembly and activation of caspase 9 and then 3 and 7. Mitochondrial permeabilization is tightly regulated by the BCL-2 family proteins that can be anti-apoptotic (BCL-2, BCL-xL or MCL-1), pro-apoptotic (BID, BIM, BAD, PUMA or NOXA) and effector pore-forming proteins such as BAX or BAK (for review see (*4*)).

Once apoptosis is triggered, this does not inevitably lead to cell death, as it is now accepted that cells can survive if mitochondrial permeabilization does not affect all mitochondria or if the ESCRT-III complex restores plasma membrane integrity (*5*) (*6*) (*7*). Currently, there is a lack of understanding of how and why the kinetics of apoptosis varies so much between cells or following different treatments.

The heterogeneous response to apoptosis is also described in isogenic cancer cell lines. This is puzzling since these cells are genetically identical and should express similar levels of both pro- and anti-apoptotic proteins. Cellular heterogeneity to undergo apoptosis thus appears to originate from non-genetic transcriptional variability. This translates into cell-to-cell variations of pro- and anti-apoptotic protein expression and activity. For instance, the Lahav group elegantly showed that there is a high variability of p53 response to DNA damaging agents; apoptotic cells accumulate p53 much faster and earlier, while this accumulation is delayed in resistant cells, allowing time for the up-regulation of pro-survival IAP (inhibitor of apoptosis) proteins (*8*).

Recent publications shed light on the effects apoptotic cells exert on neighboring cells. They can actively release EGFR ligands, FGF2 or Wnt3 to boost the survival and proliferation of neighboring cells (*9*) (*10*) (*11*). This paracrine crosstalk is also complemented by mechanotransduction mechanisms triggered by tissue stretching around apoptotic cells, which involves the Yes-associated protein (YAP) pathway (*12*).

There is therefore an increasing need to accurately estimate the variability of apoptosis induction in a given cell population. To achieve this, we developed a medium-throughput imaging pipeline using apoptotic cell markers, coupled with mathematical analysis. This allowed us to unveil a neighboring effect for apoptosis induction, irrespective of cell death stimuli or cell density, whereby the probability of undergoing apoptosis increases in the proximity of apoptotic cells. Hence, the combined application of imaging and computational analysis to evaluate the response to a given apoptotic stimulus may provide new insight into misunderstood phenomena such as fractional killing or apoptosis-induced proliferation, which are major concerns in the context of apoptosis-based cancer therapies.

## Results

### Live-cell imaging system to trigger and measure multi-stimuli apoptosis

To mathematically analyze the kinetics of apoptotic cell death while varying different parameters, we first set up a cellular system allowing medium-throughput quantification of apoptosis at the cell population level. For this, we used melanoma WM115 cells stably expressing mCherry-tagged histone H2B (H2B-mCherry) to spatially identify the exact topology of a given cell (**Figure 1A**). Following treatment with an apoptotic stimulus, the induction of apoptosis is easily visualized using the live cell-impermeant dye SYTOX Green (SG). Live cells can thus be identified by the nuclear-localized H2B-mCherry-only signal (red nuclei), whereas apoptotic cells appear yellow, owing to the colocalization of H2B-mCherry and green SG (**Figure 1A**). Next, we used various stimuli to trigger apoptosis. Treatment with TNFα and cycloheximide (CHX) was initially used as a model for death receptor-mediated apoptosis. Of note, CHX enhances TNFα-induced apoptosis by blocking the translation of short-lived anti-apoptotic proteins (*13*). Downstream of the TNFα receptor 1/2 (TNFR1/2), the DISC activates caspase 8 and triggers MOMP via BID cleavage (**Figure 1B**). To induce intrinsic apoptosis, we used two death stimuli: first, a combination of BH3 mimetics ABT-737 inactivating the anti-apoptotic proteins BCL2, BCL-xL and BCL-w, and UMI-77 targeting MCL1; second, a doxycycline (dox)-inducible WM115 cell line to activate the mitochondrial pore-forming protein BAX (**Figure 1B**). These apoptotic stimuli were then applied to SG-pre-loaded WM115 H2B-mCherry cells plated at different densities and imaged periodically (1 or 2 hours intervals) using the IncuCyte live-cell imager (**Figure 1A**).

**Figure 1.**
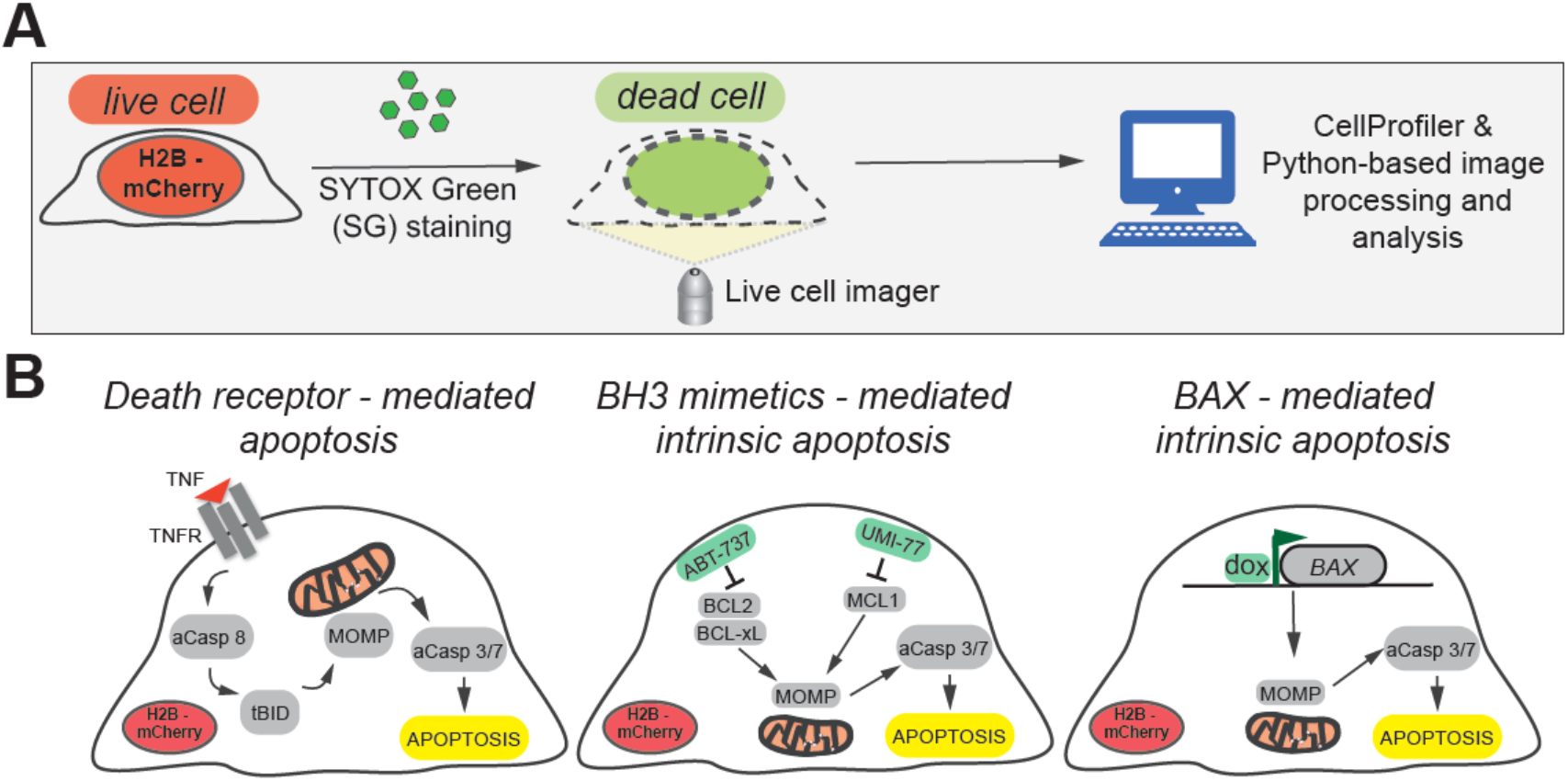
Imaging and quantification of apoptosis induction using live and dead cell markers. **A**. Overview of the live/dead cell imaging and computer analysis protocol. **B**. Apoptotic signaling induced by treatment with TNFα/CHX or BH3 mimetics, or through BAX overexpression (aCasp8 - active Caspase-8; aCasp 3/7 – active Caspase-3 and -7, tBID – truncated BID).

Next, we validated the induction of apoptosis by assessing the processing of effector caspase-3 into p17/p19 fragments and the cleavage of PARP1, which is a hallmark of apoptosis effectiveness. As shown In **Figure 2A** and **B**, treatment with TNFα/CHX (TC), BH3 mimetics (BH3m) and increasing doses of doxycycline efficiently induced apoptosis. Of note, caspase-3 processing and PARP1 cleavage were blocked by treatment with the pan-caspase inhibitor Q-VD-OPh. In addition, IncuCyte-based live-cell imaging was used to delineate the kinetics of apoptosis triggered by the different stimuli described above (**Figure 2C, D**). Apoptotic cells were marked by SG and quantified over time, with image acquisitions every 1-2 hours over a period of 48 hours on average. **Figure 2E** shows this imaging method can efficiently detect and discriminate between living cells (mCherry-positive) and apoptotic cells (SG-positive). The figure presents results of TNFα/CHX and BH3 mimetics treatment, albeit the same was observed for WM115 cells overexpressing BAX (data not shown).

**Figure 2.**
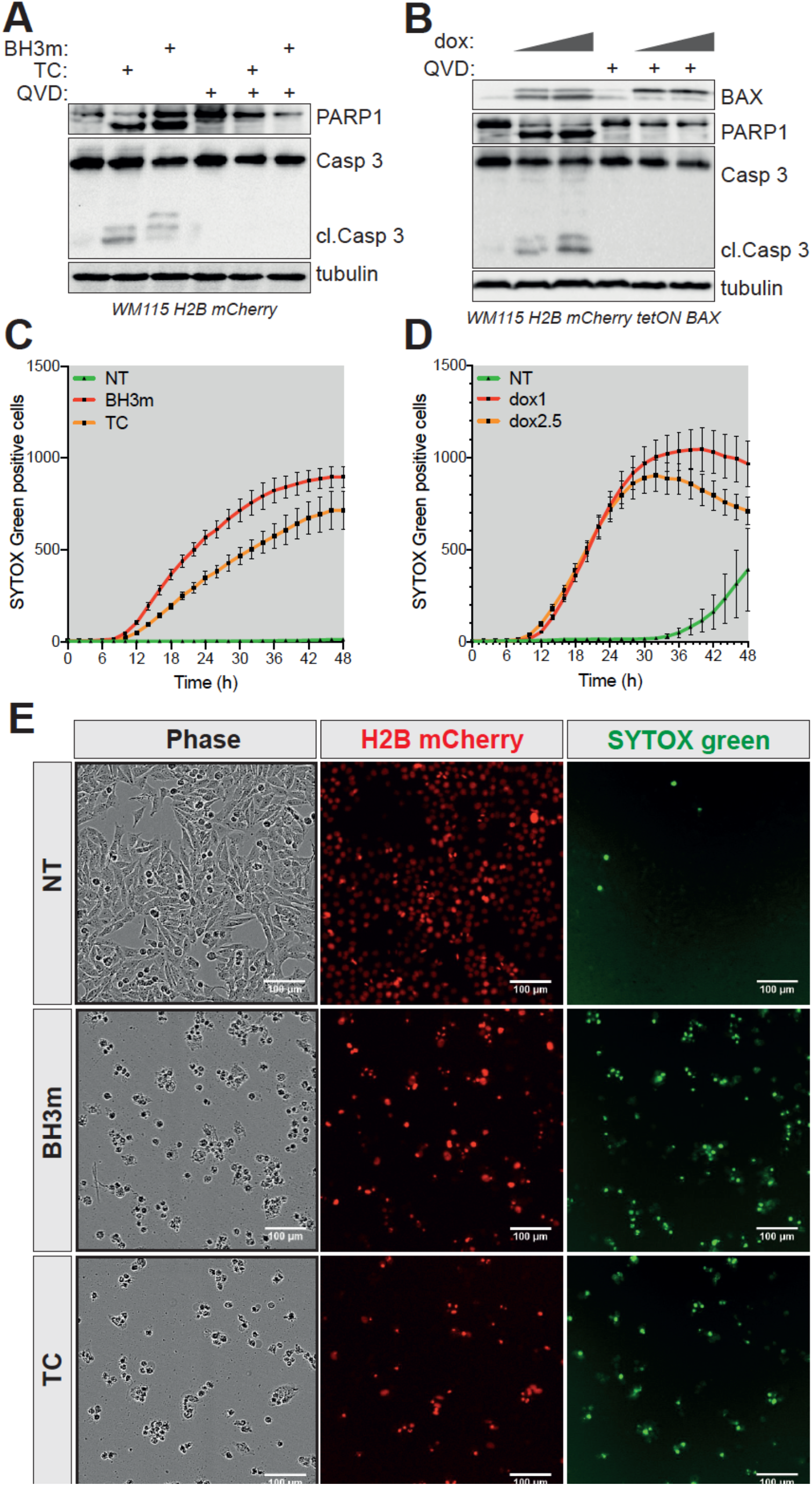
Validation of the apoptosis induction protocols. **A**. WM115 H2B-mCherry cells were treated either with TNFα/CHX (TC) (50 ng/mL of TNFα and 5 μg/mL CHX) or BH3 mimetics ABT-737/UMI-77 (10 μM each) for 12 hours in the presence or absence of the pan-caspase inhibitor Q-VD-OPh (10 μM). Protein lysates were then probed for the expression and processing of PARP1 and Caspase-3. Tubulin was used as a loading control. **B**. WM115 H2B-mCherry cells were treated with 1 or 2.5 μg/mL of doxycycline (dox) to induce BAX expression for 12 hours in the presence or absence of Q-VD-OPh (10 μM) and then analyzed as described in (A). **C-D**. IncuCyte imager-based quantification of apoptotic WM115 H2B mCherry cells (SYTOX Green-positive cells) following treatment with TNFα/CHX (TC), ABT-737/UMI-77 (**C**) or doxycycline (at 1 or 2.5 μg/mL) (**D**). **E**. Representative images of phase-contract, H2B-mCherry and SYTOX Green signal from WM115 H2B-mCherry cells treated as described in (A).

Using this cell model of apoptosis induction, coupled with medium-throughput live-cell imaging we therefore established an imaging technique able to monitor the heterogeneous induction of apoptosis at the cell population level.

### Cell clustering has no effect on the incidence of apoptosis

Following different pro-apoptotic treatments (**Figure 1B**) and IncuCyte-based live cell imaging, we obtained datasets of H2B-mCherry images (red images) and the corresponding SYTOX Green staining (green images) at different time points, for a kinetics analysis of 48 hours. We then designed two independent pipelines to segment H2B-mCherry-marked nuclei, one based on the CellProfiler software and the other written in Python, using standard image libraries to ensure a robust analysis (**Figure 1A**). The analysis pipeline included the following steps. First, for a pair of red/green images the output is a segmented “red” image with a known red object centroid position (X_*i*_, Y_*i*_) onto which a measurement of the green signal is superposed (I_*i*_, representing the maxima green signal intensity) (**Figure 3A**). Second, for a given time-point t, the analysis pipeline retrieves a set of N red centroid positions (X_*i*_, Y_*i*_) with their associated SYTOX Green intensity signal I*i*. Data analysis is then performed using three parameters δ, n and I_t_, where δ is the inter-cellular distance at which another cell(s) is/are considered to be in contact and could influence each other, n is the threshold number of neighboring cells above which a cell is considered to be in a cluster and It is the threshold intensity of SYTOX Green, derived from background noise, above which the cell is considered to be apoptotic. Of note, two cells are considered neighbors if the distance between their centroids is bellow δ (**Figure 3A**).

**Figure 3.**
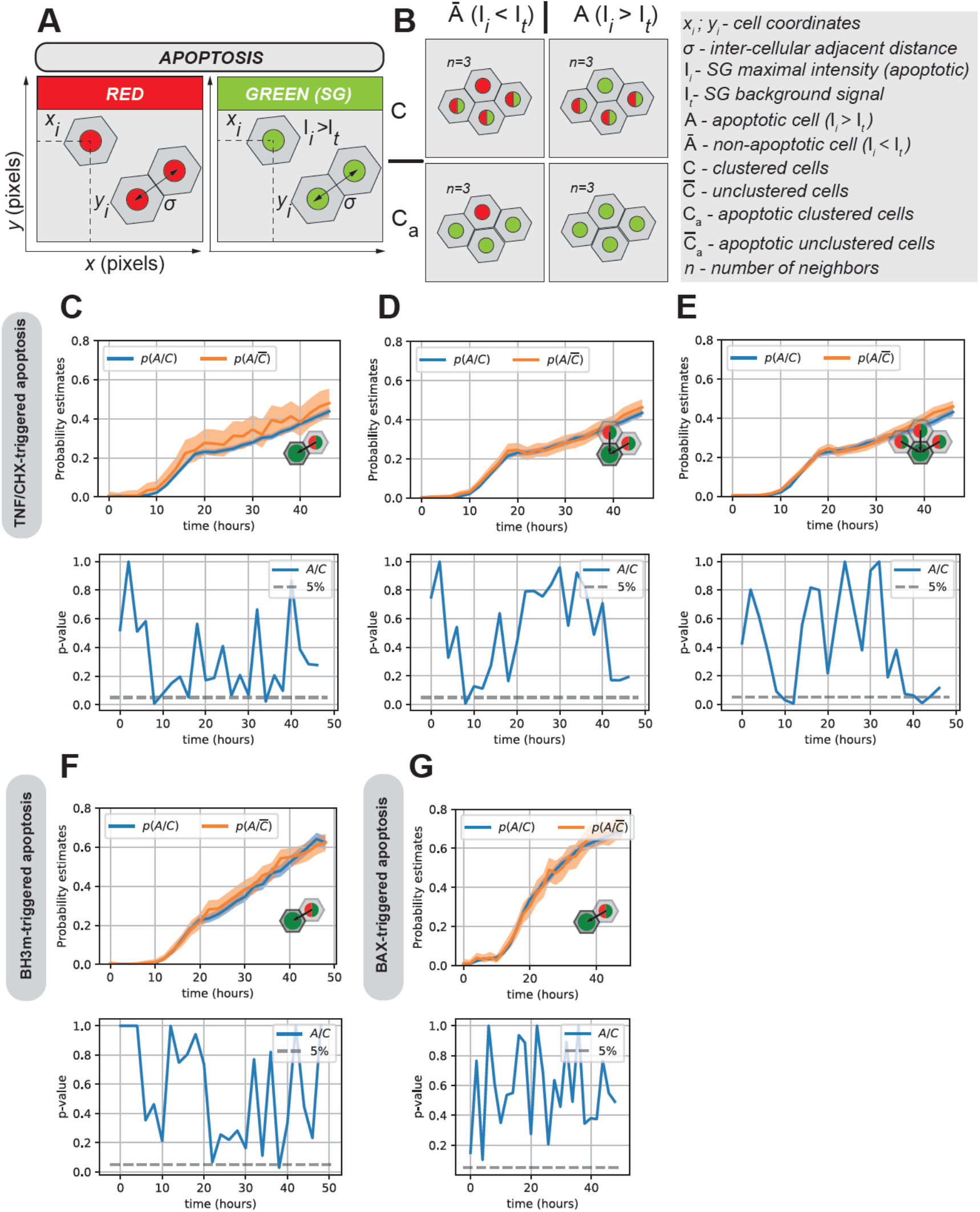
Apoptosis occurs independently of cell clustering. **A-B**. Definition of the parameters taken into account for the mathematical analysis of the effect cell clustering has on apoptosis induction. Cells with the nucleus half in red half in green are either alive or apoptotic, respectively. **C-E**. Probability estimates (upper panels) and the corresponding *p-value* (lower panels) for the occurrence of apoptosis in TNFα/CHX-treated WM115 H2B-mCherry cells surrounded by 1, 2 or 3 neighboring cells. **F-G**. Similar analysis as in (C-E), for WM115 H2B-mCherry cells treated with ABT-737/UMI-77 (**F**) and doxycycline (**G**). The neighboring effect for 1 cell is shown.

Once we determined δ, n and I_t_, we determined for each cell whether it was apoptotic (A) or not (Ā). We then established the relationships of proximity between cells (the so-called neighboring effect) by determining for each cell whether it was in a cluster of cells regardless of their apoptotic status (C) or with a known apoptotic status (Ca) (**Figure 3B**). This method is well suited to quantify spatial relations between apoptotic cells.

Using this approach, we were first interested in determining whether a correlation between cell clustering and apoptosis (between C and A). This is driven by previous work suggesting that cell density is an important modulator of both extrinsic and intrinsic apoptosis (*14*) (*15–17*).

By examining three scenarios, in which a given cell was considered to be in a cluster if it was surrounded by 1, 2 or 3 neighboring cells (regardless of their apoptotic status), we found that a cell with no neighbors 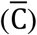 had the same probability of entering apoptosis as a cell in a cluster (of 1, 2, or 3 neighboring cells) (**Figure 3C-E** for TNFα/CHX treatment). We then computed and plotted the time dependent *p*-value of a Fisher exact test that evaluates the independence between A and C (**Figure 3C-E**, bottom panels). Of note, if the *p*-value is high (as in **Figure 3C-E**, bottom panels), this means that the hypothesis that A and C are independent cannot be rejected. In other words, this means that the probability of being apoptotic whenever one cell has neighboring relation(s) (probability p(A/C), blue curve in **Fig. 3C-E**, top panels) and the probability of being apoptotic when one cell is isolated (probability 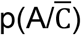, orange curve) might be equal.

The same holds true when triggering intrinsic apoptosis with BH3 mimetics (**Figure 3F**) or when overexpressing the pro-apoptotic protein BAX (**Figure 3G**). Though these results were obtained at low cell density, we confirmed their relevance for cells grown at a high density (**Supplementary Figure 1**).

Hence, these data suggest that cell clustering has no impact on the occurrence of apoptosis.

### The induction of apoptosis is enhanced by the proximity of a cell to an apoptotic cell

Next, we tested whether the proximity of a cell with one apoptotic cell or an apoptotic cluster (up to three apoptotic neighboring cells) was correlated with its propensity to undergo apoptosis. We obtained results in complete contrast to Figure 3, as the probability to enter apoptosis when in proximity with one or more apoptotic cells (probability p(A/Ca), blue curve) was significantly higher than the probability to enter apoptosis when isolated from apoptotic cells (probability p(A/C_a_), orange curve) (**Figure 4A-C** for TNFα/CHX treatment). Indeed, the *p*-value (Fisher’s exact test) was significantly low (*p* < 0.05) for the intermediate time points, meaning that both probabilities cannot be considered as equal. We observed a higher *p*-value at the beginning and the end of the kinetics, which does not affect our analysis, since there are few apoptotic cells at beginning of the time-course and, at the end, most cells are apoptotic and the neighboring effect saturates. This observation also holds true for treatment with BH3 mimetics and BAX overexpression (**Figure 4 D, E**). Finally, we validated these data at a higher cellular density (**Supplementary Figure 2**).

**Figure 4.**
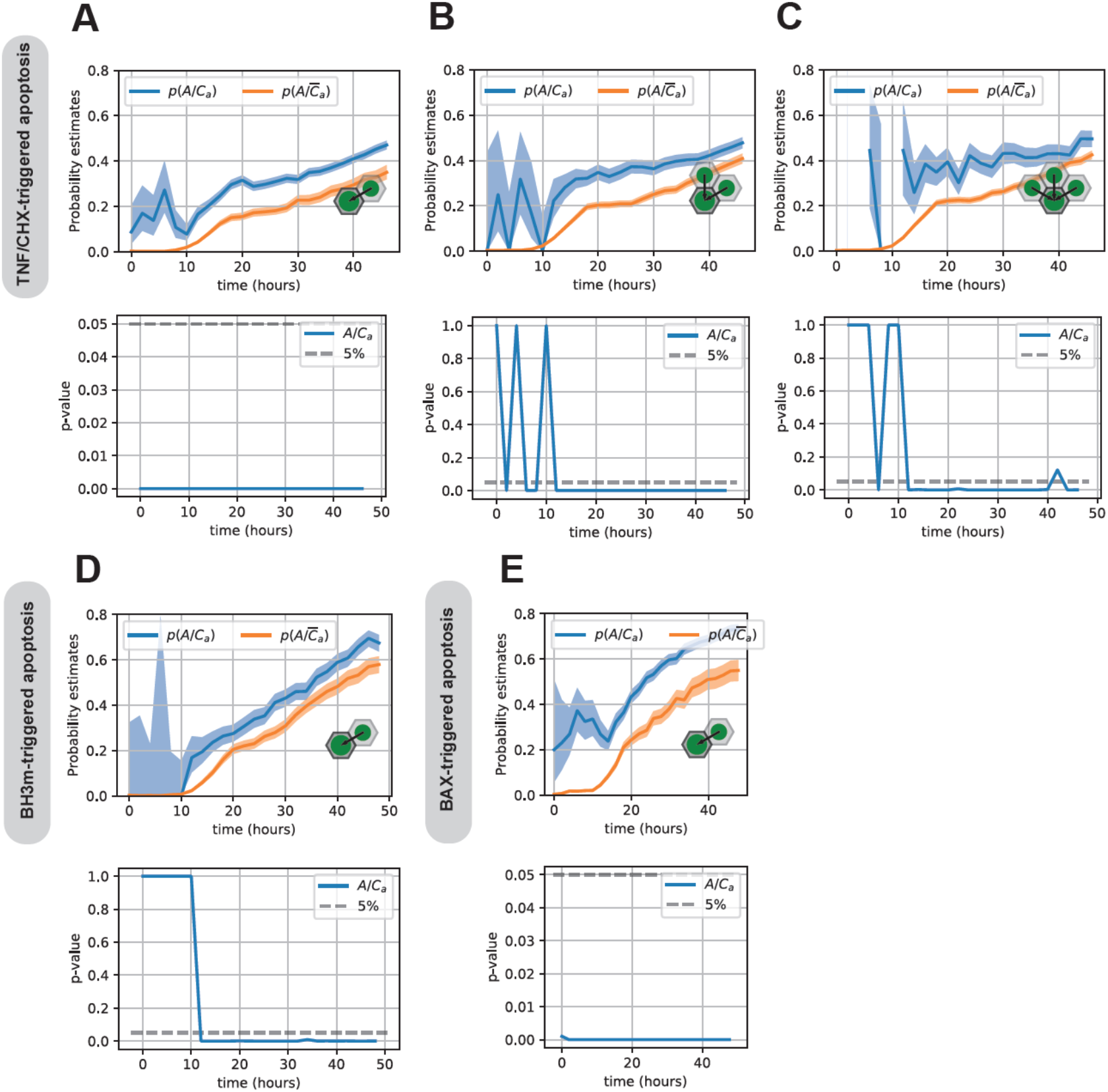
Apoptotic cell clusters influence the life-death decisions in neighboring cells. **A-C**. Probability estimates (upper panels) and the corresponding *p*-value (lower panels) for the occurrence of apoptosis in TNFα/CHX-treated WM115 H2B-mCherry cells surrounded by 1, 2 or 3 apoptotic neighboring cells. **D-E**. Similar analysis as in (A-C), for WM115 H2B-mCherry cells treated with ABT-737/UMI-77 (**D**) and doxycycline (**E**). The neighboring effect for 1 apoptotic cell is shown.

Taken together, our analysis pipeline reveals that apoptosis induction is influenced by a neighboring effect: the likelihood of a cell to undergo apoptosis is significantly higher when it is in proximity with apoptotic cells.

## Discussion

Defective induction of cell death is the cause on many diseases, ranging from cancer to neurodegenerative diseases, and it is therefore important to study its propagative nature using appropriate tools.

Here, we report the development of an analysis pipeline for medium-throughput imaging of apoptotic cancer cells that integrates several key parameters: temporal dimension, topographic localization of both living (cells stably expressing the mCherry-tagged histone 2B) and apoptotic (SYTOX Green-positive) cells, while applying several apoptotic stimuli.

Using this approach, we found that cancer cells have a similar probability of undergoing apoptosis if they are in proximity or not with other cells of unknown status. In other words, cell crowding does not have a protective effect against apoptosis. However, a cell is more likely to become apoptotic if it is in close contact with apoptotic neighboring cells. This holds true for all of the apoptotic stimuli we tested, either extrinsic or intrinsic inducers of apoptosis. These results are in line with Bhola and Simon’s study demonstrating that daughter cells, with a high likelihood of being in proximity, synchronously undergo apoptosis. This synchrony is lost as cells become less related over time (*18*). Apoptotic synchrony is therefore transiently heritable and is lost due to noise in protein translation (*19*). To take into account progeny and monitor cell proliferation history, we would need to introduce into our imaging pipeline a lineage tracer such as the one developed by Oren and colleagues (*20*).

Our experimental settings are in 2D cellular cultures, yet one might apply it to 3D cultures such as cancer organoids. For this, however, the sample will require transparization and 3D imaging microscopy. Here, the proof-of-concept is provided for apoptosis, though the protocol could easily be adapted to other types of programmed cell death, such as caspase-independent cell death, necroptosis or pyroptosis. Importantly, this method could also be helpful in identifying factors facilitating or inhibiting cell death propagation. Similar methods were developed to quantify the occurrence of fractional killing, a barrier for effective cancer treatments, and helped establish mediators such as the anti-apoptotic protein MCL1 (*21*) (*22*).

Recent research clearly demonstrates that while apoptosis is immunologically silent, it still involves releasing a plethora of signaling molecules that impinge on the fate of surrounding cells. The term of apoptosis-induced apoptosis was coined in Drosophila, where apoptotic cells in the wing disk triggered apoptosis in trans by secreting Eiger (the TNF homologue in flies), and stimulating pro-apoptotic JNK signaling (*23*). The same holds true in mammals during the regressive phase of the hair follicle (*23*). In cancer, the radiation-induced bystander effect, through which irradiated cells lead to the death of non-irradiated neighboring cells, is also a good example of this concept (*24*). Interestingly, recent studies revealed the rapid propagation of ferroptotic cell death, and the analysis pipeline described herein might provide further mechanistic insights (*25*) (*26*). Inversely, apoptosis-induced proliferation has been unveiled and describes a state in which dying cells release prostaglandins and Wnt3, instructing the proliferation of nearby cells (*27*) (*11*). Recently, the group of Tait showed that apoptotic cells also release FGF2 to enhance the anti-apoptotic resilience in trans (*10*).

The analysis method developed here could increase and enrich our knowledge on “community effects” of cell death so that they can be blocked or enhanced, according to the physio-pathological context.

## Material and methods

### Cell lines

Human melanoma cells WM115 (a gift from R. Insall – The Beatson Institute, Glasgow, UK) were maintained in DMEM supplemented with 2 mM L-glutamine (ThermoFisher Scientific, 25030-24), non-essential amino acids (ThermoFisher Scientific, 11140-035), 1 mM sodium pyruvate (ThermoFisher Scientific, 11360-039), 10% FBS (Eurobio, CVFSVF00-01) and 1% penicillin/streptomycin (ThermoFisher Scientific, 15140-122).

### Stable cell line generation

For retroviral transduction, Phoenix Ampho 293T cells (2×10^6^ in 10 cm in diameter Petri dishes) were transfected with pQXIN H2B-mCherry (gift from Dr. Luca Fava) using Lipofectamine 2000 (ThermoFisher Scientific, 11668019). Twenty-four hours later, the virus-containing supernatant was filtered and used to infect WM115 cells in the presence of 1 μg/mL polybrene (Sigma-Aldrich, H9268). This was repeated with fresh viral supernatant 48 hours later. Melanoma cells stably expressing H2B-mCherry were further purified by cell sorting. WM115 H2B mCherry cells with doxycycline-inducible expression of the BAX protein were created using the Sleeping Beauty transposon system (*28*). The plasmids used were piTR1 BAX (gift from David Goldschneider) and pCMV(CAT)T7-SB100 (Addgene, 34879) for the expression of SB100X transposase. WM115 H2B mCherry cells were then co-transfect with piTR1 BAX and pCMV(CAT)T7-SB100 plasmids using Lipofectamine 2000 and selected with puromycin (1 μg/mL).

### Western blotting

Cell lysates were prepared using NP-40 lysis buffer (1% NP-40, 1 mM EDTA, 150 mM NaCl, 50 mM Tris pH 7.4, 1 mM PMSF, Complete Protease Inhibitors (Sigma-Aldrich, 4693116001)). Protein content was determined by Bio-Rad assay, then 25-50 μg of proteins were separated on SDS-polyacrylamide gels (Biorad) under denaturating conditions (SDS PAGE sample loading buffer (VWR, GENO786-701) supplemented with 1 mM DTT) and finally transferred onto nitrocellulose. Membranes were probed with the following antibodies at a 1/1,000 dilution unless otherwise stated: BAX (2772, Cell Signaling), Caspase-3 (9662, Cell Signaling), PARP1 (9542, Cell Signaling), β-tubulin (2146, Cell Signaling).

The nitrocellulose membranes were rinsed 3 times for 10 min in TBS-Tween 0.1% and then incubated with appropriate secondary antibody coupled to HRP for 1 hour at room temperature. Finally, the proteins of interest were detected using Clarity Western ECL (Biorad, 1705060) and chemiDoc imager (Biorad, 17001401).

### Incucyte imager-based cell viability assay

Cell viability was determined using an IncuCyte Zoom imaging system (Sartorius). Cells were seeded onto Imagelock 96-well plates in media containing 30 nM SYTOX Green (Life Technologies, S7020). Following different apoptotic treatments, the cells were imaged every 60 or 120 minutes and the analysis of the SYTOX Green-positive cells was performed using the available IncuCyte image analysis software (Essen Bioscience).

### Modeling the progression of apoptosis

#### Data description

Nuclei, set as objects, are delineated at each time step using automatic segmentation of mCherry fluorescence (“Red”) images via the CellProfiler pipeline for primary objects, which is tuned to detect blob-like structures. The “apoptosis” signal is obtained by measuring the maximum intensity over each nucleus object of the “Green” fluorescence image. A threshold value for this intensity I_t_ is chosen, which defines the fluorescence intensity considered as sufficient for a given nucleus to be considered as apoptotic. In the results presented herein, the threshold chosen was 0.2 that corresponds to 2/3 to 1/2 of the maximum intensity of the Green signal.

#### Statistical test

For a given time frame, segmentation data consist in a sequence of nuclei positions (X_i_,Y_i_) and green signal intensity I_i_, with i varying from 1 to N, where N is the total number of nuclei objects for this time frame.

#### Apoptotic variable A

Given the intensity threshold I_t_, we defined the class A of apoptotic cells as the set of nuclei such that I_i_ ≥ I_t_.

#### Clustering (density) C and apoptotic clustering Ca variables

In order to separate “clustered” cells from isolated ones, we set a characteristic interaction distance δ and a threshold n on the number of neighbors. An object at position (X_i_,Y_i_) is considered to be part of a cluster (*i.e*., in the class C) if there are a least n nuclei inside the disk of radius δ and center (X_i_,Y_i_). An object at position (X_i_,Y_i_) is considered to be part of an apoptotic cluster (*i.e*., in the class C_a_) if there are a least n apoptotic nuclei (*i.e*., in the class A) inside the disk of radius δ and center (X_i_,Y_i_).

#### Independence tests

We used classical Fisher’s exact test to determine if random variables A and C are independent and subsequently if random variables A and C_a_ are independent. In this setting, for the simplicity of notation, we identify the class A with the random variable that indicates if an object belongs or not to the class A, and similarly for classes C and C_a_. We gave exact *p*-values for this test. Generally, the null hypothesis (the considered pair of random variables are independent of each other) can be rejected when the *p*-value is lower than 0.05.

## Acknowledgements

Funding from Institute Convergence PLAsCAN (ANR-17-CONV-0002), LabEx DEVweCAN (University of Lyon), Agence Nationale de la Recherche (ANR) Young Researchers Project (ANR-18-CE13-0005-01), La Ligue Nationale Contre le Cancer and Fondation de France supported this work. We thank Brigitte Manship for reviewing the manuscript.

## Author contributions

<u>Conceptualization</u>, G. Ichim, D. Coulette, H.L. and E.G.; <u>Methodology</u>, G. Ichim, D. Coulette, H.L. and E.G.; <u>Formal analysis</u>, G. Ichim, D. Coulette and H.L.; <u>Investigation</u>, G. Ichim, D.C., D. Coulette and H.L.;<u>Resources</u>, G. Ichim, D. Coulette, H.L. and E.G.; <u>Writing</u> – Original Draft and Editing, G. Ichim; All authors reviewed and edited the manuscript; <u>Supervision</u>, G. Ichim, D. Coulette, H.L. and E.G.; <u>Project administration and funding acquisition</u>, G. Ichim.

**Supplementary Figure 1 (related to Figure 3).**
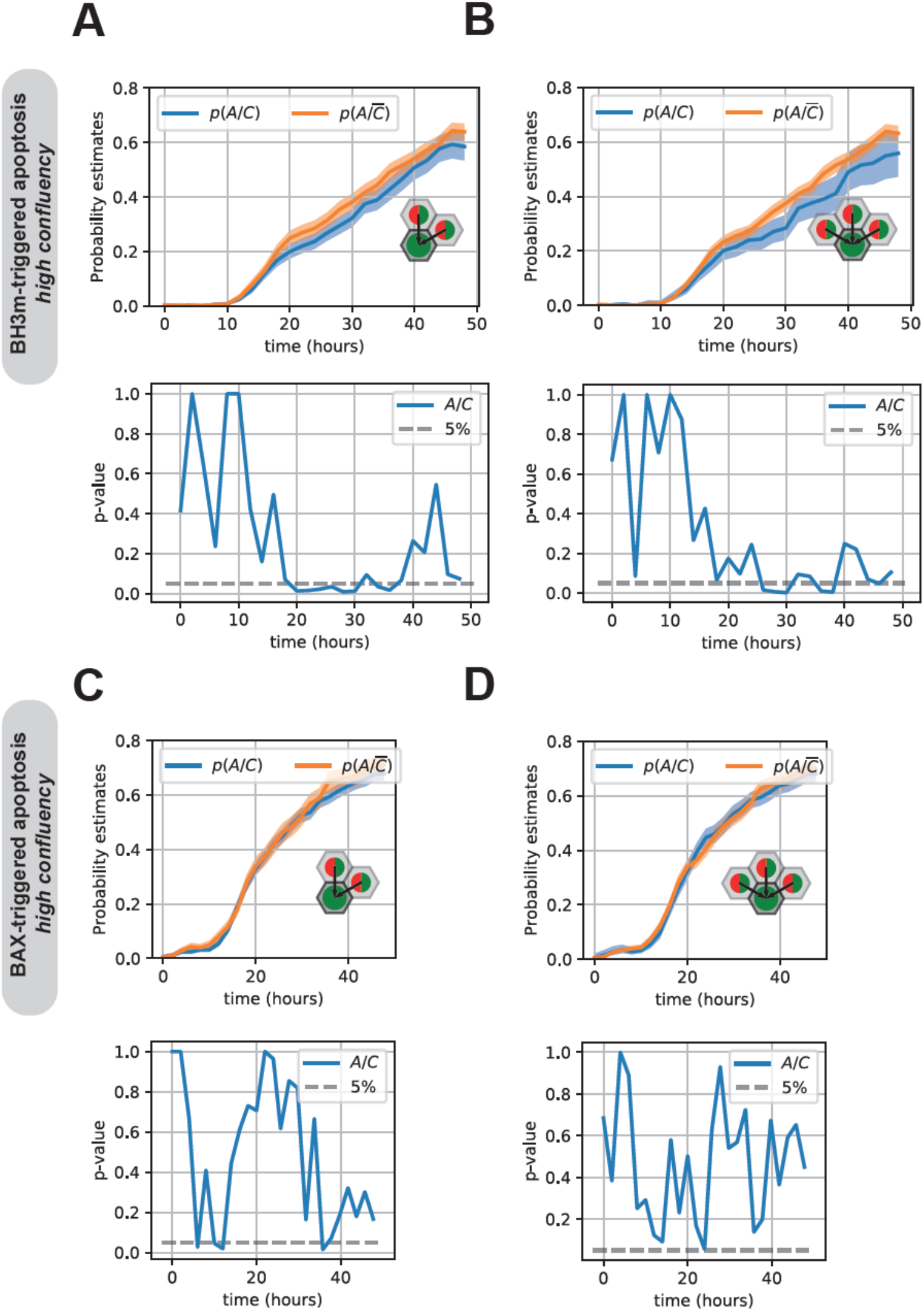
**A, B**. Probability estimates (upper panels) and the corresponding *p*-value (lower panels) for the occurrence of apoptosis in ABT-737/UMI-77-treated WM115 H2B-mCherry cells surrounded by 2 or 3 neighboring cells. **C, D**. Similar analysis as in (A, B), albeit apoptosis was triggered by doxycycline induction of pro-apoptotic BAX protein.

**Supplementary Figure 2 (related to Figure 4).**
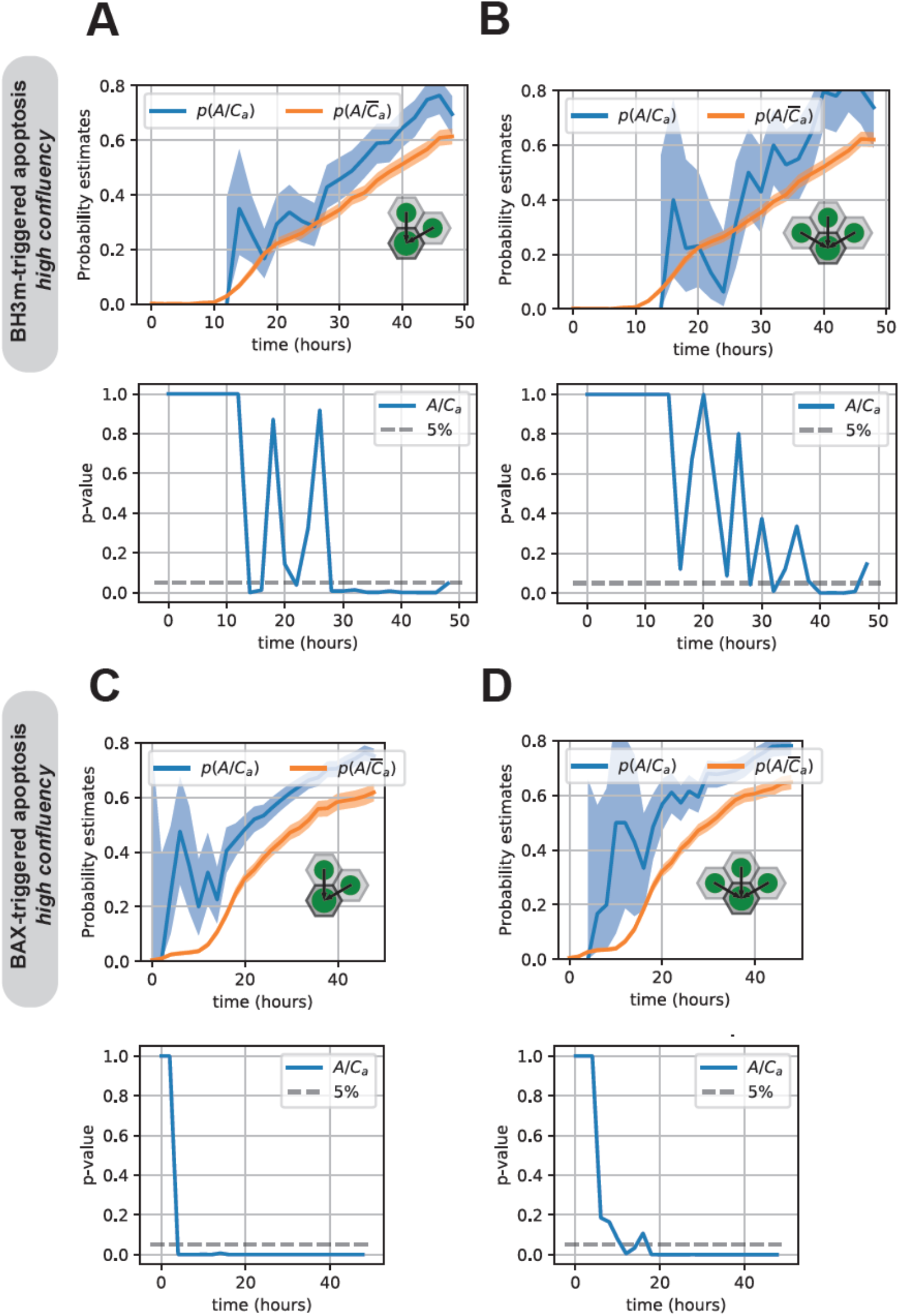
**A, B**. Probability estimates (upper panels) and the corresponding *p*-value (lower panels) for the occurrence of apoptosis in ABT-737/UMI-77-treated WM115 H2B-mCherry cells surrounded by 2 or 3 apoptotic neighboring cells. **C, D**. Similar analysis as in (A, B), albeit apoptosis was triggered by doxycycline induction of pro-apoptotic BAX protein.

